# The contribution of genetic and environmental effects to Bergmann’s rule and Allen’s rule in house mice

**DOI:** 10.1101/2021.06.14.448454

**Authors:** Mallory A. Ballinger, Michael W. Nachman

**Affiliations:** Department of Integrative Biology and Museum of Vertebrate Zoology, University of California, Berkeley, Berkeley, CA 94702-3160

**Keywords:** body size, extremity length, adaptive plasticity, heritability, *Mus*

## Abstract

Distinguishing between genetic, environmental, and genotype-by-environment effects is central to understanding geographic variation in phenotypic clines. Two of the best-documented phenotypic clines are Bergmann’s rule and Allen’s rule, which describe larger body sizes and shortened extremities in colder climates, respectively. Although numerous studies have found inter- and intraspecific evidence for both ecogeographic patterns, we still have a poor understanding of the extent to which these patterns are driven by genetics, environment, or both. Here, we measured the genetic and environmental contributions to Bergmann’s rule and Allen’s rule across introduced populations of house mice (*Mus musculus domesticus*) in the Americas. First, we documented clines for body mass, tail length, and ear length in natural populations, and found that these conform to both Bergmann’s rule and Allen’s rule. We then raised descendants of wild-caught mice in the lab and showed that these differences persisted in a common environment and are heritable, indicating that they have a genetic basis. Finally, using a full-sib design, we reared mice under warm and cold conditions. We found very little plasticity associated with body size, suggesting that Bergmann’s rule has been shaped by strong directional selection in house mice. However, extremities showed considerable plasticity, as both tails and ears grew shorter in cold environments. These results indicate that adaptive phenotypic plasticity as well as genetic changes underlie major patterns of clinal variation in house mice and likely facilitated their rapid expansion into new environments across the Americas.

## Introduction

Clines in phenotypes have historically been attributed to natural selection, reflecting adaptation to local environments (Huxley 1939; Endler 1977). Two of the best described clinal patterns in animals are Allen’s rule and Bergmann’s rule. Allen’s rule is the observation that extremities, such as limb length and tail length, are shorter in colder climates compared to warmer regions, resulting in latitudinal clines (Allen 1877). Bergmann’s rule is the observation that body sizes are larger in colder climates, resulting in latitudinal clines in body size (Bergmann 1847). Shortened extremities and larger body sizes minimize heat loss by reducing surface area to volume ratios and are thus viewed as thermoregulatory adaptations (Mayr 1956). Numerous studies have documented Bergmann’s rule and Allen’s rule within and across species of birds (Johnston and Selander 1964; James 1970; Laiolo and Rolando 2001; Romano et al. 2020) and mammals (Brown and Lee 1969; Griffing 1974; Yom-Tov and Nix 1986; Fooden and Albrecht 1999), including humans (Ruff 1994; Ruff 2002; Foster and Collard 2013; Betti et al. 2015). Moreover, various meta-analyses have supported the generality of these rules (Ashton et al. 2000; Ashton 2002; Freckleton et al. 2003; Meiri and Dayan 2003; Blackburn and Hawkins 2004; Millien et al. 2006; Nudds and Oswald 2007; Olson et al. 2009; Symonds and Tattersall 2010; Alhajeri et al. 2020). On the other hand, several meta-analyses have questioned the ubiquity of these patterns arguing that statistical support is weak (Geist 1987; Gohli and Voje 2016; Riemer et al. 2018) or that phenotypic differences are more likely to be driven by resource abundance or other factors rather than by considerations of temperature (Scholander 1955; McNab 1971; Geist 1987; Alhajeri and Steppan 2016; Alroy 2019). The contradicting results found across the literature are unsurprising given the variation within and among datasets, such as choice of taxonomic groups, environmental variables, and inconsistencies in measurements. Moreover, virtually all studies to date are based on observations of individuals sampled in natural populations in which factors such as age, reproductive condition, social status, pathogen and parasite loads, and overall health are not easily controlled. Thus, we still have very little understanding of the mechanisms underlying Allen’s rule and Bergmann’s rule.

Missing from many of these discussions are careful analyses determining which traits are genetically encoded, environmentally influenced, or both. Environmentally influenced traits are phenotypically plastic traits, and these traits may also harbor genetic variation for plasticity (i.e., genotype-by-environment interactions) (Des Marais et al. 2013). Most traits associated with Bergmann’s rule and Allen’s rule are complex, meaning they are both polygenic and strongly influenced by the environment (Falconer and Mackay 1996; Lynch et al. 1998; Yang et al. 2010; Harpak and Przeworski 2021). In fact, in his original description of clinal patterns, Allen (1877) emphasized the role of the environment in directly modulating phenotypes. Disentangling genetic from non-genetic effects in natural populations is difficult when using phenotypic data collected from wild-caught animals. Genetic contributions to trait values may be masked by environmental effects (plasticity) and genotype-by-environment interactions (Conover and Schultz 1995; Alho et al. 2011). Phenotypic plasticity may also generate clinal patterns, giving a false impression of adaptive clines (James 1983). In fact, many temporal changes in body size are driven by the environment and not genetic adaptation in birds (Teplitsky et al. 2008; Husby et al. 2011) and mammals (Ozgul et al. 2009, 2010). Furthermore, we have little understanding of how populations conforming to these ecogeographic rules vary in the degree and direction of plasticity they exhibit in response to environmental stimuli. Variation in plasticity (i.e., genotype-by-environment interactions) may facilitate adaptation and divergence in polygenic traits (Via and Lande 1985; Gillespie and Turelli 1989; Gomulkiewicz and Kirkpatrick 1992; West-Eberhard 2003). However, controlling for environmental effects and measuring the contributions of phenotypic plasticity is difficult, as transplant experiments and common garden experiments are infeasible for many taxa. These limitations have impeded our ability to make substantial progress on understanding the evolutionary and ecological mechanisms underlying Bergmann’s rule and Allen’s rule.

House mice (*Mus musculus domesticus*) provide a tractable system for disentangling the genetic and environmental contributions to complex traits. House mice have recently expanded their range from Western Europe to the Americas, where they can be found from the tip of South America to Alaska. Across this broad latitudinal range, mice are exposed to various environmental gradients, including both temperature and aridity (Phifer-Rixey and Nachman 2015). Despite residing in these novel environments for only a few hundred generations, there is evidence for clinal variation across latitudes. Specifically, mice in eastern North America follow Bergmann’s rule (Lynch 1992; Phifer-Rixey et al. 2018), with larger mice in more northern populations. These body size differences persist in a common environment over several generations, indicating that they have a genetic basis (Lynch 1992; Phifer-Rixey et al. 2018).

Selection on house mice over ten generations in the laboratory recapitulates these clinal patterns: mice bred at lower temperatures become larger and undergo genetic divergence in body size (Barnett and Dickson 1984). Furthermore, previous work has revealed an environmental influence on tail length when exposed to cold temperatures. Specifically, house mice reared in a cold environment grew significantly shorter tails than mice reared at warm temperatures, consistent with Allen’s rule (Sumner 1909, 1915; Barnett 1965). However, these earlier studies investigated only a single population of mice or used classical inbred laboratory strains of mice, making it difficult to place the results in an explicit evolutionary framework. We still have little understanding of the phenotypic variation of house mice across their entire latitudinal distribution, and even less understanding of the contributions of genetics and environment to these complex traits.

Here, we use a combination of approaches to tease apart genetics from plasticity in Bergmann’s rule and Allen’s rule in house mice from North and South America. First, we determined if house mice conform to both Bergmann’s rule and Allen’s rule across North and South America by analyzing phenotypic data from wild-caught individuals. Second, we collected individuals from temperate and tropical populations of house mice from the ends of their latitudinal distribution, brought them back to the lab, and established wild-derived colonies. We analyzed phenotypic differences between populations and across generations in a common lab environment and identified a heritable basis to both Allen’s rule and Bergmann’s rule. Third, to measure the influence of environment on body size and extremity length, we performed a second common garden experiment by rearing both lab populations of mice using a full-sib design in a cold and warm environment and measured the effects on body size and extremity length. Measuring developmental plasticity within and between populations allowed us to assess the influence of temperature on complex traits and to understand the evolutionary mechanisms underlying these clinal patterns. Specifically, we found that unlike body size, tail and ear length are highly plastic and this plastic response goes in the same direction as the evolved response of New York mice, highlighting an example of adaptive phenotypic plasticity.

## Materials and Methods

### Phenotypic data from wild-caught mice

To determine if house mice conform to Allen’s rule and Bergmann’s rule, we tested for associations between body mass, tail length, ear length, and latitude in wild house mice collected across North and South America. We downloaded specimen data of all house mouse records from VertNet (Constable et al. 2010) on October 13, 2020, using the search query: *vntype*:specimen, *genus*:Mus. We obtained 62,139 museum records and retained records that included *Mus musculus* specimens collected in North or South America (excluding islands). We omitted individuals explicitly listed as pregnant, juvenile, subadult, or immature, and included individuals listed as adult, mature, or with no age class or reproductive condition specified. We also manually coded females and males as ‘ adult’if they fulfilled any of the following criteria: females - presence of placental scars, parous, or lactating; males - presence of seminal vesicles, testes descended (TD), or testes scrotal (TS). Tail lengths shorter than 20mm and longer than 120mm (*n* = 8), and ear lengths greater than 30mm (*n* = 1) were considered extreme outliers (greater than 3.5 standard deviations from the mean) and were removed from downstream analyses. Sample information for the final VertNet dataset (*n* = 3,018) is provided in Data S1.

We assessed the overall relationship between body mass, extremity length, and latitude by fitting linear models in R (v. 4.1.1), including sex as a covariate (Table S1). We tested for clinal patterns of body mass and extremity length across latitude in wild-caught house mice using Spearman correlations for each sex separately. To test if mice from colder regions conform to Bergmann’s rule and Allen’s rule, we extracted mean annual temperature data for all VertNet sampling locations at 1 km (30-seconds) spatial resolution from WorldClim 2.1 (Fick and Hijmans 2017). We tested for clinal patterns of body mass and extremity length across mean annual temperature (°C) using Spearman correlations for each sex separately.

### Laboratory-reared mice - common garden experiment 1

To disentangle genetic effects from environmental effects, we collected live animals from two locations that represent the ends of a latitudinal transect: Manaus, Amazonas, Brazil (MAN), located near the equator at 3°S latitude, and Saratoga Springs, New York, USA (SAR), located at 43°N latitude. Details of this common garden experiment are given in (Phifer-Rixey et al. 2018) and (Suzuki et al. 2020). Briefly, live mice from both Brazil (*n* = 38) and New York (*n* = 30) were brought back to the lab at the University of California, Berkeley. Within each population, unrelated pairs of wild-caught mice were mated to produce first generation (N1) lab-reared mice. Mice were then paired in sib-sib matings to generate inbred lines. These inbred lines (New York: *n* = 10 lines; Brazil: *n* = 12 lines) have been maintained through sib-sib matings for over 10 generations. Wild-caught mice and their descendants were housed in a standard laboratory environment at 21°C with a 12-hr dark and 12-hr light cycle. Water and commercial rodent chow (Teklad Global, 18% protein, 6% fat) were provided ad libitium. Standard museum measurements (total length, tail length, hind foot length, ear length, and body mass) were taken for all wild-caught and inbred mice from each population (see Data S2). We removed outliers for tail length (< 50mm (*n* = 2)) and ear length (< 8mm (*n* = 1)) from downstream analyses.

### Estimating heritability

To estimate heritability for body mass and extremity length in house mice, we performed midparent-offspring regression on N2 (parents) and N3 (offspring) laboratory-born mice. Generations N2 and N3 were chosen to eliminate any residual environmental and maternal effects that could influence heritability estimates. Midparent values were calculated as the mean trait value between mother and father, and heritabilities were calculated as the regression coefficients (slopes) of offspring values against midparent values (Lynch et al. 1998). We performed midparent-offspring regressions on 13 Brazil families (representing 10 different inbred lines) and 13 New York families (representing 10 different inbred lines). We also calculated regression coefficients of offspring values against maternal and paternal values separately for all three traits. Heritabilities from these regressions were estimated as twice the slope of the regression of offspring values against maternal or paternal values (Lynch et al. 1998). We assessed the significance of regression coefficients for each heritability estimate using ANOVAs, implemented in the car package (v. 3.0.11) (Fox and Weisberg 2019).

### Developmental phenotypic plasticity - common garden experiment 2

To determine the influence of phenotypic plasticity on body mass and extremity length, we performed a second common garden experiment by rearing laboratory-born mice from both populations in a cold and warm environment. We used temperature as the environmental variable because temperature is highly correlated with latitude (Millien et al. 2006) and phenotypic variation in wild house mice across North and South America is explained most by temperature-related variables (Suzuki et al. 2020). Specifically, we used two wild-derived inbred lines each from Brazil (MANA, MANB) and New York (SARA, SARB). Each line has been inbred for more than 10 generations, and thus mice within a line harbor reduced levels of genetic variation. Roughly equal numbers of males and females were produced for each within-line comparison (New York: *n* = 40; Brazil: *n* = 40; see Data S3 and S4). These population-specific sample sizes align with previous experimental studies in house mice (e.g., Phifer-Rixey et al. 2018). Full-sibs were born at room temperature (21°C) and singly-housed at weaning (∼21 days old). After a brief acclimation period, we randomly assigned 3.5-week-old mice into size-matched groups based on sex-specific body mass, and then housed mice at either 5°C or remained at 21°C for the duration of the experiment (∼50 days total). We measured initial body mass and tail length and recorded subsequent body mass and tail lengths once a week for each mouse. At the end of the experiment, we euthanized mice at 75 ± 3 days of age, and recorded final body mass and tail length, in addition to standard museum measurements. Two final ear lengths were not included in downstream analyses due to ear damage. We deposited skulls and skeletons of all mice in the Museum of Vertebrate Zoology, University of California, Berkeley (catalog numbers are given in Data S4). All experimental procedures were in accordance with the UC Berkeley Institutional Animal Care and Use Committee (AUP-2017-08-10248).

### Data Analysis

All data analyses and visualizations were completed in R (v. 4.1.1). Within R, we used the tidyverse (v. 1.3.1) (Wickham et al. 2019), performance (v. 0.8.0) (Lüdecke et al. 2021), cowplot (v. 1.1.1), here (v. 1.0.1), and rmarkdown (v. 2.11) (Allaire et al. 2021) packages, along with R base library. Relative tail length and relative ear length were calculated by dividing tail or ear length by body mass for each individual. We also performed all analyses using tail length residuals and ear length residuals (by regressing length from body mass across individuals) and obtained similar results.

For common garden experiments 1 and 2, we fitted linear mixed models in the lme4 package (v. 1.1.27.1) (Bates et al. 2015) to determine if morphology varied between sex, population, generation, or environment. In each model, we included: 1) the morphological trait as the response variable; 2) sex, population, generation (experiment 1) or environment (experiment 2), and their interaction as fixed effects; and 3) inbred lines as a random effect. The significance of interactions was evaluated using ANOVA based on type III (partial) sums of squares, implemented in the car package (v. 3.0.11) (Fox and Weisberg 2019). We performed *post hoc* comparisons on significant two-way interactions using Tukey’s HSD tests (*P* < 0.05). Lastly, we calculated the effect size (ω2) of predictors using the effectsize package (v. 0.5) (Ben-Shachar et al. 2020) to evaluate the relative influence of sex, population, and generation or environment on body mass and extremity length (Olejnik and Algina 2003) (Table 1; Table S2).

**Table 1.** Results of linear mixed models investigating the effects of sex, population, environment, and their interaction on body mass and extremity length in house mice.

| Trait | Factor | $\chi^2$ | DF | P-value | Effect size ($\omega^2$) |
| --- | --- | --- | --- | --- | --- |
| Body Mass (g) |  |  |  |  |  |
|  | Sex | 42.15 | 1 | < 0.001 | 0.36 |
|  | Population | 4.00 | 1 | 0.045 | 0.43 |
|  | Environment | 1.72 | 1 | 0.19 | <0.01 |
| Relative Tail Length (mm/g) |  |  |  |  |  |
|  | Sex | 42.44 | 1 | < 0.001 | 0.37 |
|  | Population | 6.54 | 1 | 0.011 | 0.58 |
|  | Environment | 25.59 | 1 | < 0.001 | 0.25 |
|  | Population:Environment | 3.71 | 1 | 0.054 | 0.04 |
| Relative Ear Length (mm/g) |  |  |  |  |  |
|  | Sex | 45.62 | 1 | < 0.001 | 0.39 |
|  | Population | 3.72 | 1 | 0.054 | 0.40 |
|  | Environment | 20.07 | 1 | < 0.001 | 0.21 |

## Results

### Evidence for Bergmann’s rule and Allen’s rule in wild house mice

We assessed the relationship between tail length, ear length, body mass, and latitude in mice collected across North and South America to determine if populations of house mice conform to Allen’s rule and Bergmann’s rule. Using a large dataset downloaded from VertNet (*n* = 3,018; Data S1), we found little evidence for Bergmann’s rule, as body mass showed a non-significant, positive correlation with latitude across both males and females (Figure 1A). In contrast, we found stronger evidence for Allen’s rule in house mice from the Americas, with both tail length (Figure 1C) and ear length (Figure 1E) showing a significant, negative correlation with latitude. These patterns of extremity length largely hold true across both sexes (Figure 1C, 1E). In each of these comparisons, however, latitude explained less than 1% of the phenotypic variation (Table S1). Similar patterns of Bergmann’s rule and Allen’s rule were seen using mean annual temperature (°C) as a predictor (Figure S1). Specifically, body mass declined and extremity length increased with increasing mean annual temperature (Figure S1A,C,E).

**Figure 1.**
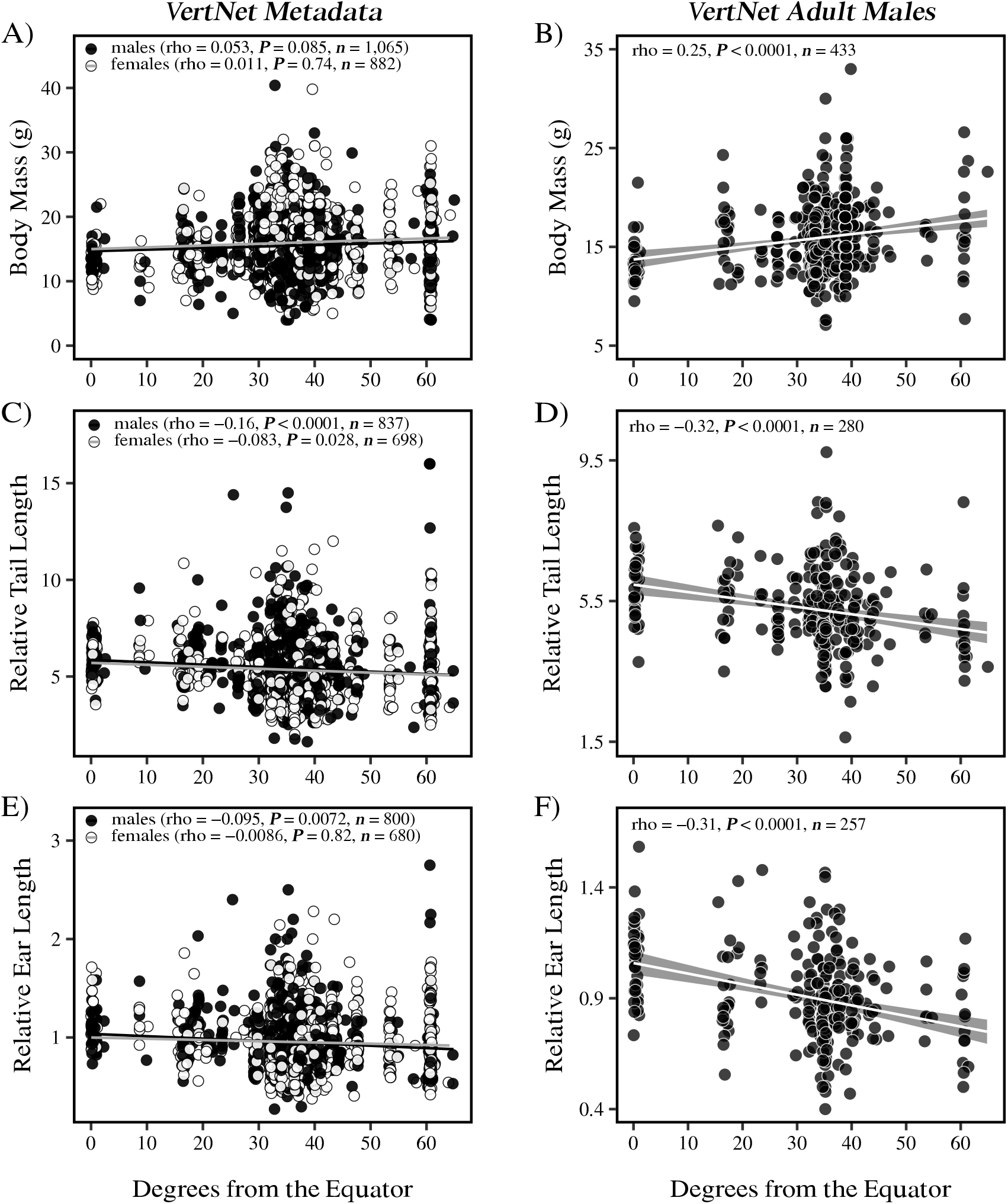
Bergmann’s rule and Allen’s rule in house mice from North and South America. Associations between body mass (A-B), tail length (C-D), ear length (E-F), and absolute latitude across wild-caught North and South American house mice. Tail length and ear length are plotted relative to body mass for each individual. Individuals are represented as individual points, with males depicted in black and females depicted in white. Results from Spearman correlations are presented in each plot, along with sample sizes. For clarity, standard error shading is omitted from linear regression lines associated with the VertNet Metadata (panels A, C, and E).

The lack of evidence for Bergmann’s rule in Figure 1A and S1A may be due to the influence of uncontrolled factors (e.g., age, diet, health), environmental effects, or phenotypic plasticity. Although we minimized variation in age and reproductive status by removing explicitly labeled pregnant females, juveniles, and subadults, we still see large variation across all three traits, likely due to various factors that were not recorded. To reduce this variation, we further filtered the VertNet dataset to only include explicitly labeled adult males (Figure 1B, 1D, 1F; *n* = 445). We focused on males since females show more variation in body mass than males (Levene’s test; *F* = 17.89; *P* = 0.005), likely due to reproductive condition. In this more controlled set of adult males, we see strong evidence for both Bergmann’s rule (Figures 1B, S1B) and Allen’s rule (Figures 1D, 1F, S1D, S1F). The comparison between the larger dataset and the more curated dataset highlights how uncontrolled variation in collated museum metadata may obscure broad ecogeographic patterns.

### Differences in body mass and extremity length persist in a common environment and are heritable

The phenotypic clines observed across wild house mice could represent genetic differences, phenotypic plasticity, or both. To disentangle genetics from plasticity, we collected live mice from near the equator (Manaus, Amazonas, Brazil) and from 43°N latitude (Saratoga Springs, New York, USA) and brought them into a common laboratory environment. Population-specific differences in body mass and extremity length in wild-caught mice (N0) persisted across generations of laboratory-reared mice (Figure S2). Specifically, mice from New York were larger than mice from Brazil (ANOVA, *P* < 0.001, ω^2^= 0.56) (Figure S2A; Table S2). New York mice also had shorter tails (ANOVA, *P* < 0.001, ω^2^= 0.65) (Figure S2B; Table S2) and shorter ears (ANOVA, *P* < 0.001, ω^2^= 0.58) (Figure S2C; Table S2) compared to mice from Brazil. The maintenance of body mass and extremity length differences in a common environment and across generations suggests that these traits are heritable in house mice.

To estimate the heritability (*h2*) of body mass and extremity length in New York and Brazil house mice, we performed midparent-offspring regressions on N2 and N3 mice (Figure 2). Both Brazil and New York mice yielded similar, high heritability values for body mass and extremity length. Specifically, Brazil mice showed significant and very similar heritabilities for all three traits (ANOVA, *P* < 0.05; Figure 2). These heritability estimates are in general agreement with previous estimates of body mass and tail length in house mice (Rutledge et al. 1973). Furthermore, using single-parent offspring regressions, both New York and Brazil mice showed similar heritabilities for all three traits (Figure S3). Maternal- and paternal-offspring regressions suggest that body mass and extremity length in New York and Brazil mice are not determined by inheritance of just one sex, since heritability estimates are in rough agreement with midparent-offspring regression estimates. Overall, these results suggest that body mass and extremity length are heritable and under strong genetic control in house mice.

**Figure 2.**
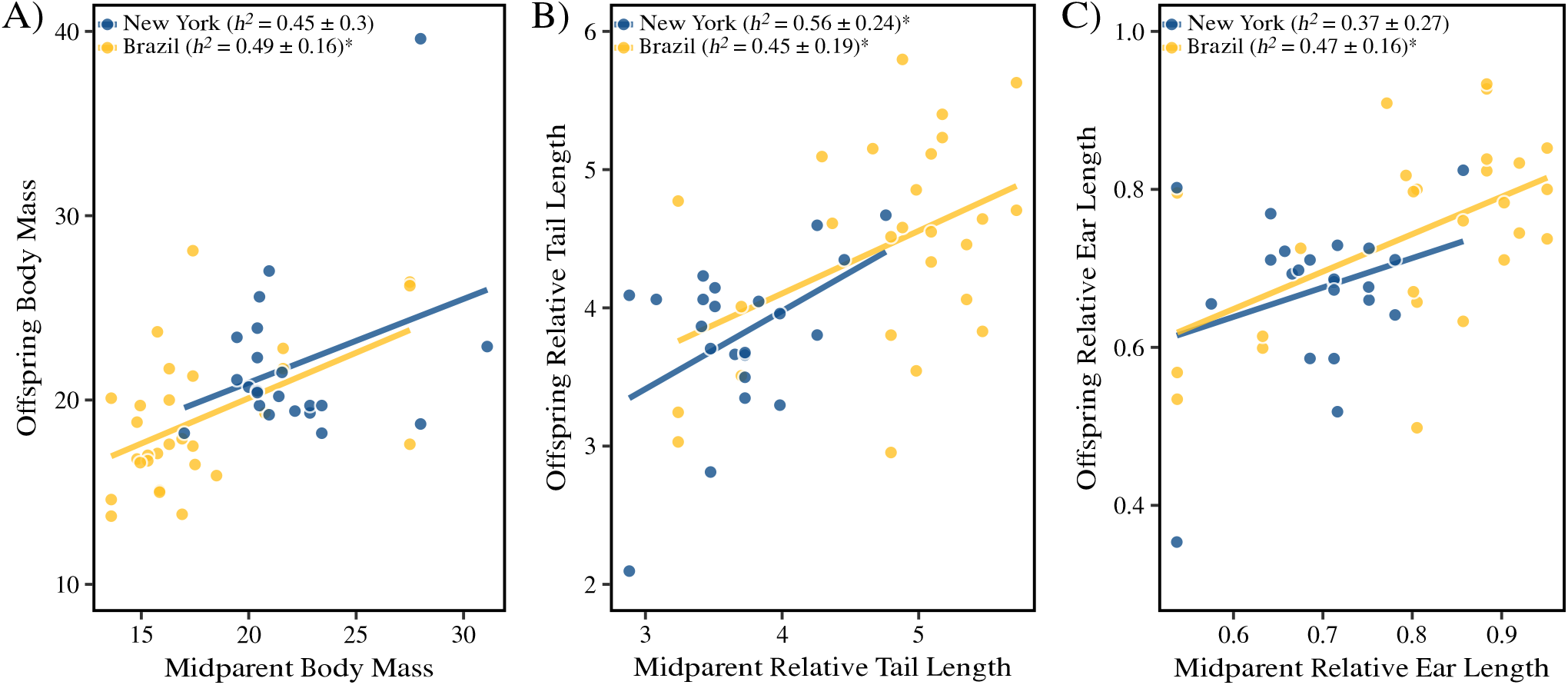
Heritability (*h*^*2*^) estimates for body mass and extremity length in New York and Brazil house mice. Midparent-offspring regressions were performed on body mass (A), relative tail length (B), and relative ear length (C). Midparent values were calculated as the mean trait value between mothers and fathers for 13 families in each Brazil and New York populations. Heritabilities (*h*^*2*^ ± standard error) were estimated as slopes of midparent-offspring regressions. Significance of each regression was assessed with ANOVAs and is represented with an asterisk. Sample sizes: New York: *n* = 22; Brazil: *n* = 26.

### Extremity length, but not body size, is greatly influenced by temperature

The results presented above identified phenotypic divergence in body mass and tail and ear length in house mice, with New York mice having shorter tails and ears and larger body sizes than mice from Brazil, consistent with Allen’s rule and Bergmann’s rule, respectively. To determine the influence of phenotypic plasticity on these traits, we reared laboratory-born mice from both populations in a cold and warm environment. Genetic differences in body mass between inbred lines of New York and Brazil mice were evident at weaning (Figure 3A), with New York mice larger than Brazil mice. These body mass differences between populations persisted across developmental stages from 3 to 11 weeks. In all cases, males were larger than females. Full-sibs (of the same sex) reared at different temperatures showed no differences in body mass (Figure 3A). At the end of the experiment, body size differences recapitulated patterns seen across generations, with New York mice larger than Brazil mice (Table 1; Figure 3B) and males larger than females (Table 1; Figure 3B). The lack of plasticity in body mass was not a result of differences in fat accumulation, as body mass index (BMI) did not differ between populations (ANOVA, χ^2^=0.50, *P*>0.05) or environments (ANOVA, χ^2^=1.28, *P*>0.05) (Figure S4). These results suggest that phenotypic plasticity does not play a significant role in body size evolution of house mice.

**Figure 3.**
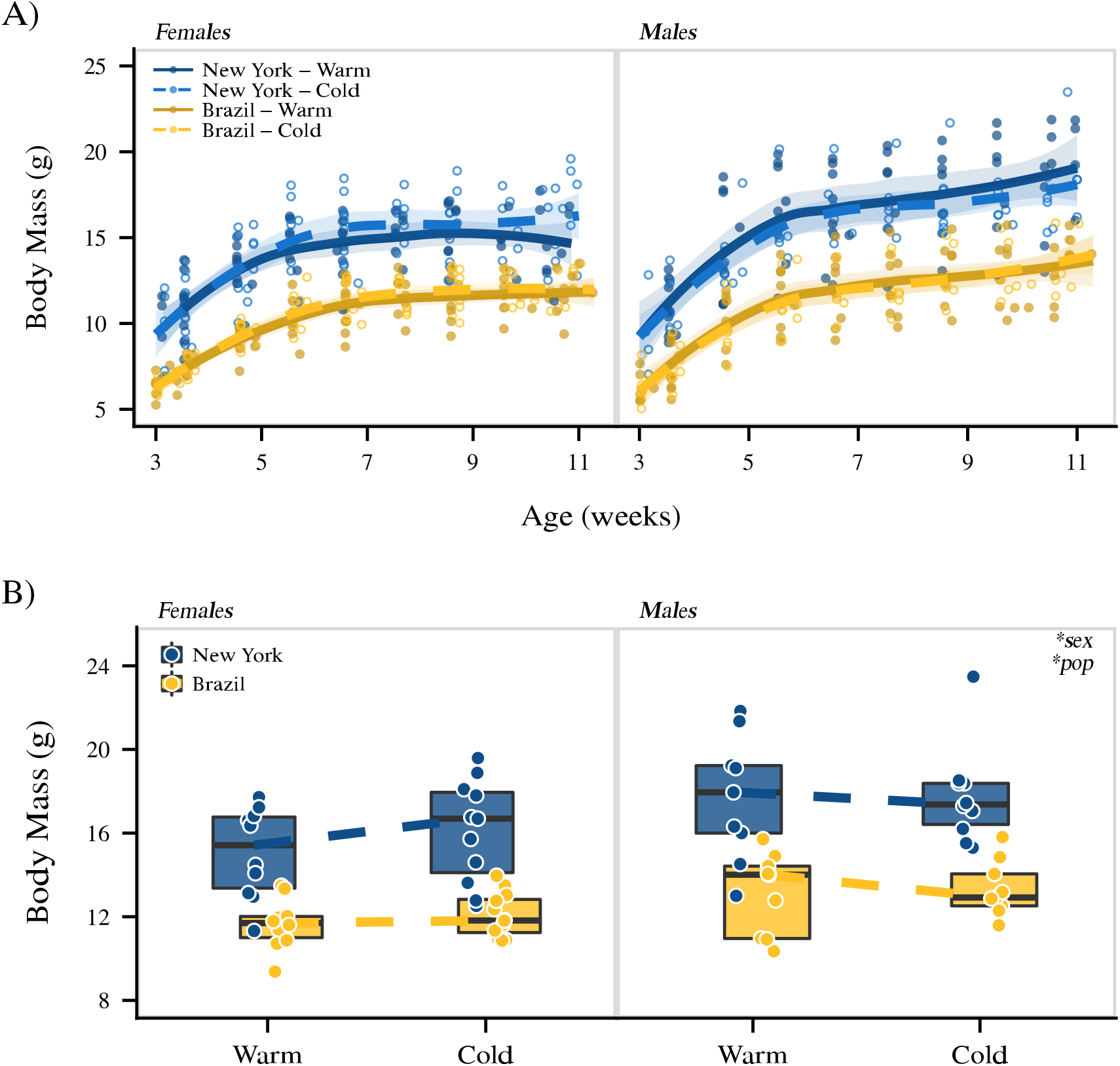
Genetic differences and very little plasticity in body mass among New York and Brazil house mice. A) Body mass growth trajectories across environments and 11 weeks of development in New York (blue) and Brazil (gold) house mice. Individuals are plotted as individual points, with cold-reared mice denoted as open circles and warm-reared mice denoted as filled circles (New York: *n* = 20 per treatment; Brazil: *n* = 20 per treatment). Population means are depicted as smoothed regression fits, with cold mice denoted as dashed lines and warm mice denoted as solid lines. Each line is accompanied with standard error shading. B) Individuals at 11 weeks of age are represented as individual points (New York: *n* = 20 per treatment; Brazil: *n* = 20 per treatment), and boxplots indicate the 25th, median, and 75th quartiles. Dashed lines connect median values of each genotype across warm and cold environments. Results from linear mixed models are presented in the upper right corner (*P<0.05; Table 1).

In contrast to body mass, tail length was greatly influenced by developmental temperature, with the first few weeks post-weaning having the greatest influence on absolute tail length (Figure 4). Specifically, inbred mice reared in a cold environment grew shorter tails than inbred mice reared in a warm environment (Table 1; Figures 4, 5A). The magnitude of this effect was striking in Brazil mice, corresponding to 5.40 mm, on average, or 7% of the total adult tail length. Despite developmental temperature playing a significant role in tail length, genetic differences in tail length were evident at the end of the experiment, with Brazil mice growing longer tails than New York mice in both environments (Table 1; Figures 4, 5A). Thus, tail length exhibits both genetic divergence and phenotypic plasticity. Similarly, cold-reared mice grew shorter ears than warm-reared mice (Table 1; Figure 5B), and the magnitude of the plastic response corresponded to 5% of the total adult ear length in both populations. These temperature-growth responses of the extremities were not a simple consequence of body size differences, as body mass did not differ between treatments (Figure 3). Overall, unlike body size, extremity length showed significant plasticity in response to temperature, with mice growing shorter extremities in a cold environment, consistent with patterns of Allen’s rule.

**Figure 4.**
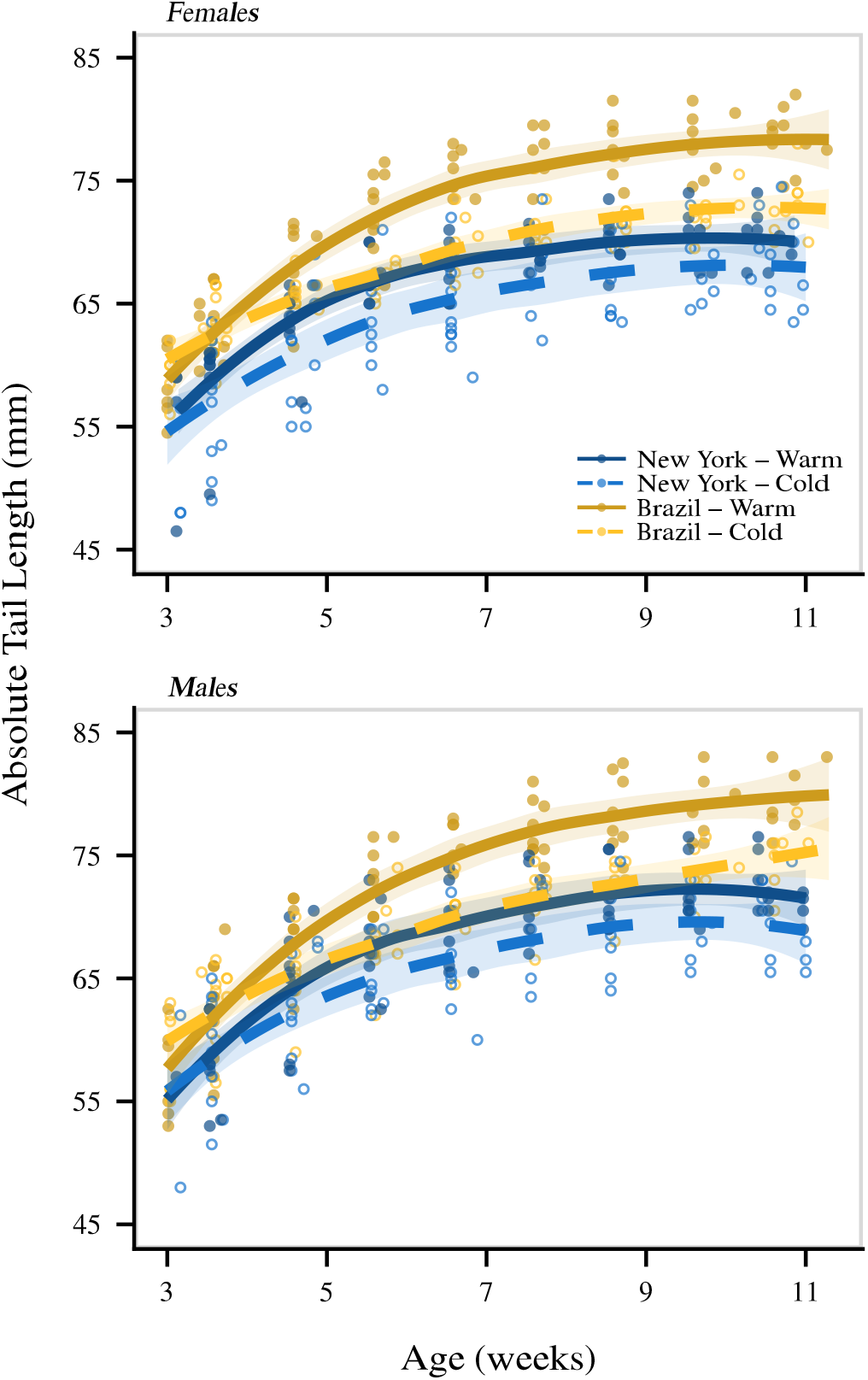
Tail length is highly influenced by cold temperature across development. Absolute tail length growth trajectories across environments in New York (blue) and Brazil (gold) house mice. Individuals are plotted as individual points, with cold-reared mice denoted as open circles and warm-reared mice denoted as filled circles (New York: *n* = 20 per treatment; Brazil: *n* = 20 per treatment). Population means are depicted as smoothed regression fits, with cold mice denoted as dashed lines and warm mice are denoted as solid lines. Each line is accompanied with standard error shading. The same individuals depicted here are also depicted in Figure 5.

**Figure 5.**
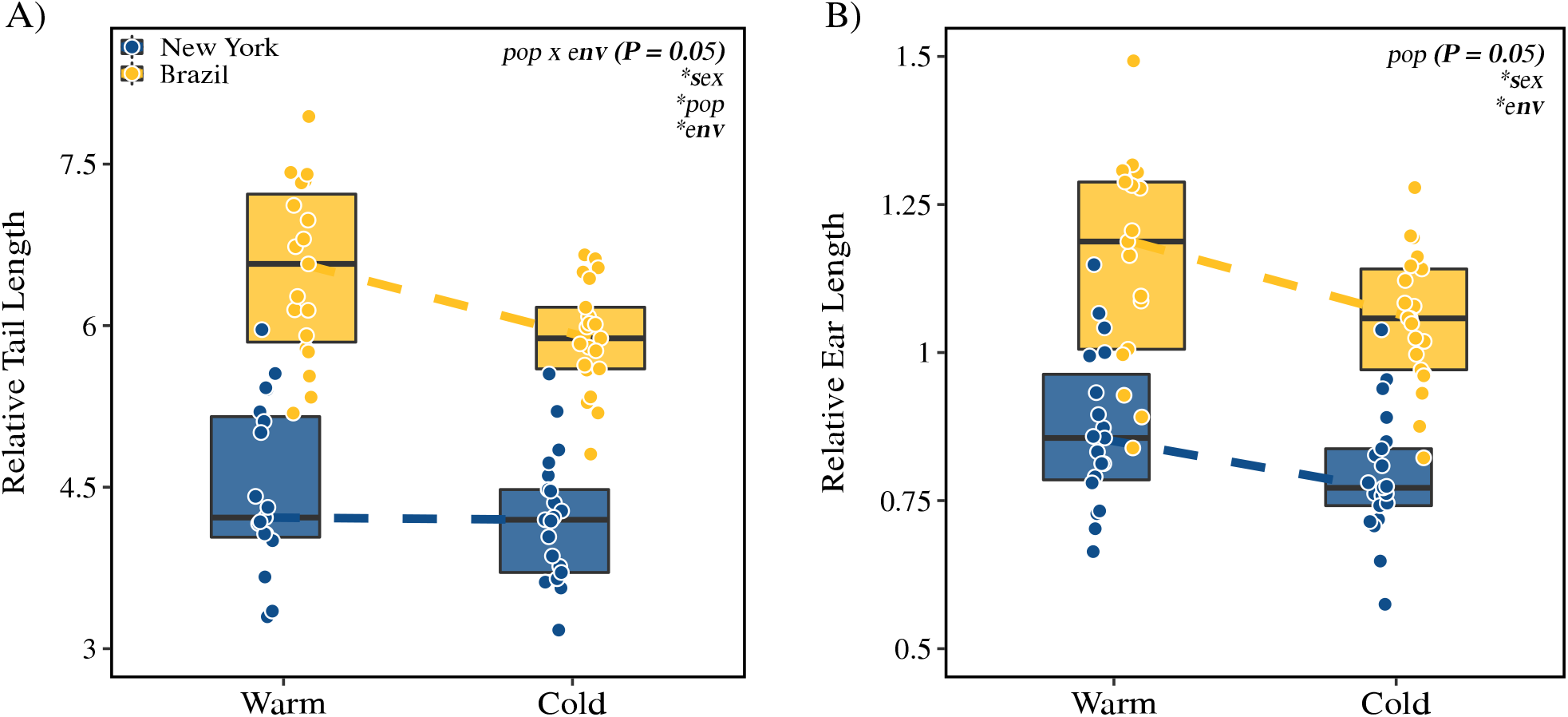
Adaptive phenotypic plasticity in extremity length among New York and Brazil house mice. Tail length (A) and ear length (B) are plotted relative to body mass for each individual. Individuals are represented as individual points (A) New York: *n* = 20 per treatment; Brazil: *n* = 20 per treatment; B) New York: *n* = 20 per treatment; Brazil: *n* = 19 per treatment). Boxplots indicate the 25th, median, and 75th quartiles. Dashed lines connect median values of each genotype across warm and cold environments. Both sexes were combined for simplicity. Results from linear mixed models are presented in the upper right corner (\**P*<0.05; Table 1). The same individuals depicted here are also depicted in Figure 4.

### Adaptive phenotypic plasticity in extremity length

Differences between warm- and cold-reared mice revealed a strong plastic response to temperature in extremity length. Because plasticity is considered adaptive when a phenotype is altered in the same direction as natural selection (Ghalambor et al. 2007), we next asked whether phenotypic plasticity of Brazil mice goes in the same or opposite direction as the evolved response of New York mice. For absolute tail length, the mean trait value of Brazil house mice reared in the cold nearly recapitulates the tail length of New York mice reared in a warm environment (Figure 4), highlighting an example of adaptive phenotypic plasticity. Interestingly, the plastic response of tail length for New York mice (i.e., difference between warm New York tail length and cold New York tail length) was attenuated in comparison to the plastic response of tail length for Brazil mice (Figures 4, 5A), suggesting that New York house mice may be closer to the phenotypic optimum or that there is a developmental constraint on minimum tail length. Lastly, plasticity in ear length of Brazil house mice went in the same direction as the evolved response of New York house mice reared in a warm environment (Figure 5B), further illustrating adaptive phenotypic plasticity. The overall degree and direction of plasticity in extremities mirror patterns associated with Allen’s rule. In fact, the plastic response of both tail length and ear length in Brazil mice (i.e., difference between warm Brazil trait value and cold Brazil trait value) explains roughly 40% of the mean phenotypic differences we observe in wild mice (i.e., between N0 Brazil mice and N0 New York mice).

## Discussion

Previous studies have provided conflicting assessments of the generality of Bergmann’s and Allen’s rules and have rarely combined field and laboratory studies to identify the contribution of genetic and non-genetic effects to phenotypic variation. Here, we focused on one species that has recently expanded its range across many degrees of latitude, and we studied phenotypic variation in the wild and in the lab. Moreover, by rearing inbred mice at different temperatures we were able to assess the contribution of phenotypic plasticity to patterns seen in nature.

First, we found that wild house mice across North and South America conform to Bergmann’s rule and Allen’s rule, as house mice are larger in size with shortened extremities farther from the equator and at colder temperatures. Second, heritable differences in body mass and tail and ear length in a common environment indicated a genetic basis to Bergmann’s rule and Allen’s rule, presumably reflecting thermoregulatory adaptations. Finally, we measured the contributions of phenotypic plasticity to these traits and found that tail and ear length are highly plastic in response to cold temperature, while body size is not. The plastic response in extremity length to cold temperatures appears adaptive, matching the direction of change of extremity length seen in wild, temperate house mice. Adaptive plasticity associated with Allen’s rule, in conjunction with strong selection for body size, likely promoted the rapid expansion of house mice into new environments across the Americas.

### Genetic contributions to ecogeographic rules

Parallel phenotypic clines across multiple transects provide strong evidence for natural selection (Endler 1977). Body mass in house mice increases as latitude increases (Figure 1A-B), resembling patterns seen in eastern North America (Lynch 1992; Phifer-Rixey et al. 2018), South America (Suzuki et al. 2020), and Australia (Tomlinson and Withers 2009). These clinal patterns are consistent with Bergmann’s rule. Notably, this pattern is much clearer when looking at a dataset that includes only adult males (Figure 1B) compared to a dataset with all animals (Figure 1A). This difference may help to explain the discrepancies between previous meta-analyses based on museum collections (e.g., Ashton et al. 2000; Meiri and Dayan 2003; Riemer et al. 2018). Moreover, the substantial trait variation in Figure 1 among individuals, even when sampled at the same latitude, provides strong motivation for studying these traits in a common laboratory environment.

Phenotypic measurements of lab-reared mice from temperate and tropical populations revealed a heritable basis for the difference in body mass, with mice from a colder climate significantly larger than mice from a tropical environment (Figure 2A). This pattern was seen across generations (Figure S2) and heritability estimates for body mass were relatively high across both populations (Figure 2). These results agree with previous studies that also found a genetic basis for body size differences in house mice from eastern North America (Lynch 1992; Phifer-Rixey et al. 2018), western North America (Ferris et al. 2021), and South America (Suzuki et al. 2020), and suggest that there has been strong directional selection for body size in house mice. Selection over short time scales leading to latitudinal clines for body size has also been shown for other non-native species, such as in the genus *Drosophila*. Specifically, body size clines in introduced species of *Drosophila* have been repeated across continents, in common garden experiments, and through experimental evolution studies (Cavicchi et al. 1985; Coyne and Beecham 1987; Partridge et al. 1994; James et al. 1995; Land et al. 1999; Huey et al. 2000; Gilchrist et al. 2001, 2004). Latitudinal clines for body size have also arisen rapidly in introduced populations of house sparrows (Johnston and Selander 1964, 1971) and European starlings (Cardilini et al. 2016). Together, these results suggest that introduced species may undergo strong selection while colonizing new environments, allowing patterns conforming to Bergmann’s rule to become quickly established.

Phenotypic measurements of wild mice revealed clines for relative tail length and relative ear length, with shorter extremities seen in mice collected farther from the equator and at colder temperatures, consistent with Allen’s rule (Figures 1, S1). As with body size, a clearer picture of clinal variation emerged when only considering adult males (Figures 1D, 1F, S1D, S1F) compared to all animals. Measurements of lab mice showed that these differences are heritable (Figures 2, S2). Shorter tails in northern populations of house mice may minimize heat loss and thus be an adaptation to the cold, as tail length shows a positive correlation with temperature of the coldest month across rodents (Alhajeri et al. 2020). Similar trends and correlations have also been found for limb length and bill length in birds (Nudds and Oswald 2007; Symonds and Tattersall 2010; Danner and Greenberg 2015; Friedman et al. 2017). In addition to thermoregulatory advantages, alternative mechanisms for Allen’s rule have been postulated to explain why longer tails are found in the tropics, such as enhanced climbing ability with increased arboreality (Alroy 2019; Mincer and Russo 2020). Because house mice are commensal with humans, it is unlikely that longer tails confer a climbing advantage in the tropics.

### Contributions of phenotypic plasticity to ecogeographic rules

Body size in house mice shows very little plasticity in response to cold temperature (Figure 3), reaffirming that there has likely been strong directional selection for body size in house mice. Lack of plasticity associated with Bergmann’s rule is consistent with previous studies in laboratory mice (Sumner 1909, 1915; Ashoub 1958; Serrat et al. 2008; Serrat 2013) and, in addition to selection, may be due to a number of physiological factors. The environmental influence of temperature may need to occur pre-weaning or prenatal to elicit a plastic response (e.g., Weaver and Ingram 1969; Burness et al. 2013; Andrew et al. 2017). Exposure to high temperatures instead of low temperatures may elicit a plastic response in body size, as seen previously in some endotherms (Ashoub 1958; Gordon 2012; Burness et al. 2013; Andrew et al. 2017).

Unlike Bergmann’s rule, Allen’s rule can be generated via developmental phenotypic plasticity, as extremity length is highly sensitive to ambient temperature in both mammals and birds (Serrat 2014; Tattersall et al. 2017). Our results in house mice agree with previous studies in mammals, with mice growing shorter tails and ears in a cold environment (Figure 5) (Ogle and Mills 1933; Harland 1960; Chevillard et al. 1963; Weaver and Ingram 1969). In laboratory mice, temperature directly affects the growth of cartilage in both tails and ears, influencing extremity length (Serrat et al. 2008). Furthermore, the widespread patterns of tail length plasticity in response to cold are also recapitulated at the skeletal level, with both the length and number of caudal vertebrae decreasing in response to cold temperatures in mice (Barnett 1965; Noel and Wright 1970; Thorington Jr 1970; Al-Hilli and Wright 1983). Although we did not measure skeletal differences between New York and Brazil mice, it seems likely that the tail length plasticity we observed is a result of plasticity in both number and length of individual caudal vertebrae. Moreover, ear length shows the greatest plasticity in both populations, with both New York and Brazil mice growing shorter ears in the cold. The pronounced plastic response of ears compared to tails may indicate that smaller appendages consisting entirely of cartilage are less developmentally canalized. Less constraint associated with extremities may also underly the highly plastic nature of Allen’s rule compared to Bergmann’s rule. This is illustrated by tail length and ear length plasticity accounting for roughly 40% of the observed differences among wild New York and Brazil house mice.

### Adaptive phenotypic plasticity and Allen’s rule

Phenotypic plasticity is adaptive when it aligns with the direction of selection, moving traits closer to the local phenotypic optima (Baldwin 1896; West-Eberhard 2003; Ghalambor et al. 2007). We found evidence for adaptive phenotypic plasticity underlying Allen’s rule, as plasticity produced shorter ears and tails in cold environments. We also observed an attenuated plastic response for tail length in New York house mice compared to Brazil house mice, suggesting that New York mice are closer to the phenotypic optimum and are better adapted to colder environments. Overall, plasticity in house mouse extremities mirrors general evolutionary patterns of shorter extremity lengths in colder climates and may play an important role in generating Allen’s rule.

There are two ways by which adaptive phenotypic plasticity can facilitate the colonization of new environments. Adaptive plasticity can incompletely move the trait value closer to the phenotypic optimum, with directional selection refining the trait value, leading to subsequent genetic changes (Price et al. 2003; Ghalambor et al. 2007). Alternatively, adaptive plasticity can slow or impede evolution by moving individuals completely to the phenotypic optimum, shielding genetic variation from natural selection (Price et al. 2003; Ghalambor et al. 2007). We find evidence for the first scenario for both tail and ear length in house mice.

Specifically, both genetic and plastic contributions generate shorter tails and ears in colder environments. Despite the plastic response of extremity length in Brazil mice explaining roughly 40% of the mean phenotypic differences observed in wild mice, we see clear evidence of genetic differences in tail length and ear length between New York and Brazil house mice. This suggests that phenotypic plasticity moves extremity length close to the local optimum but does not shield it from subsequent selection. Overall, adaptive phenotypic plasticity in addition to strong, directional selection underlying Bergmann’s rule, likely facilitated the rapid expansion of house mice into new environments across the Americas.

## Acknowledgements

We thank Michael Sheehan and Felipe Martins for collecting wild mice, and we thank Kathleen Ferris, Gabriela Heyer, Dana Lin, Felipe Martins, Megan Phifer-Rixey, Michael Sheehan, and Taichi Suzuki for help with mouse husbandry. We also thank Jesse Alston, Libby Beckman, Sylvia Durkin, Emilie Richards, Michelle St. John, Molly Womack, Daniel Bolnick, David Lowry, and two anonymous reviewers for their constructive feedback that improved the manuscript. Funding and support for this work was provided by the National Institutes of Health (R01 GM074245 and R01 GM127468), an American Society of Mammalogists Grant-in-Aid of Research, graduate student research funds from the Museum of Vertebrate Zoology and Department of Integrative Biology, a National Science Foundation Graduate Research Fellowship (DGE-1106400), Junea W. Kelly Museum of Vertebrate Zoology Graduate Fellowship, and a UC Berkeley Philomathia Graduate Fellowship.

## Statement of Authorship

Both authors conceived and designed the study and obtained funding. M.A.B. performed data collection, analysis, and visualization with contributions and guidance from M.W.N. The manuscript was written by M.A.B. and edited by M.W.N.

## Data and Code Availability

The code to perform analyses for this study is available as a git-based version control repository on GitHub (https://github.com/malballinger/Ballinger_allenbergmann_AmNat_2021). The analysis can be reproduced using a GNU Make-based workflow with built-in bash tools (v. 3.81) and R (v. 4.1.1). Supporting data files and scripts have been deposited on Zenodo (https://zenodo.org/record/5823597).

## Supplemental Tables

**Table S1.** Results of linear models investigating the relationship between body mass, extremity length, and absolute latitude. Sex was included as a covariate in each model.

| Trait | Adj. R<sup>2</sup> | Factor | F-value | DF | P-value | Effect size ($\omega^2$) |
| --- | --- | --- | --- | --- | --- | --- |
| Body Mass (g) | 0.008 |  |  |  |  |  |
|  |  | Absolute Latitude | 11.53 | 1 | < 0.001 | <0.001 |
|  |  | Sex | 4.74 | 1 | 0.03 | <0.001 |
| Relative Tail Length (mm/g) | 0.009 |  |  |  |  |  |
|  |  | Absolute Latitude | 14.62 | 1 | <0.001 | <0.001 |
|  |  | Sex | 0.89 | 1 | 0.35 | <0.001 |
| Relative Ear Length (mm/g) | 0.005 |  |  |  |  |  |
|  |  | Absolute Latitude | 9.21 | 1 | 0.002 | <0.001 |
|  |  | Sex | 0.08 | 1 | 0.78 | <0.001 |

**Table S2.**
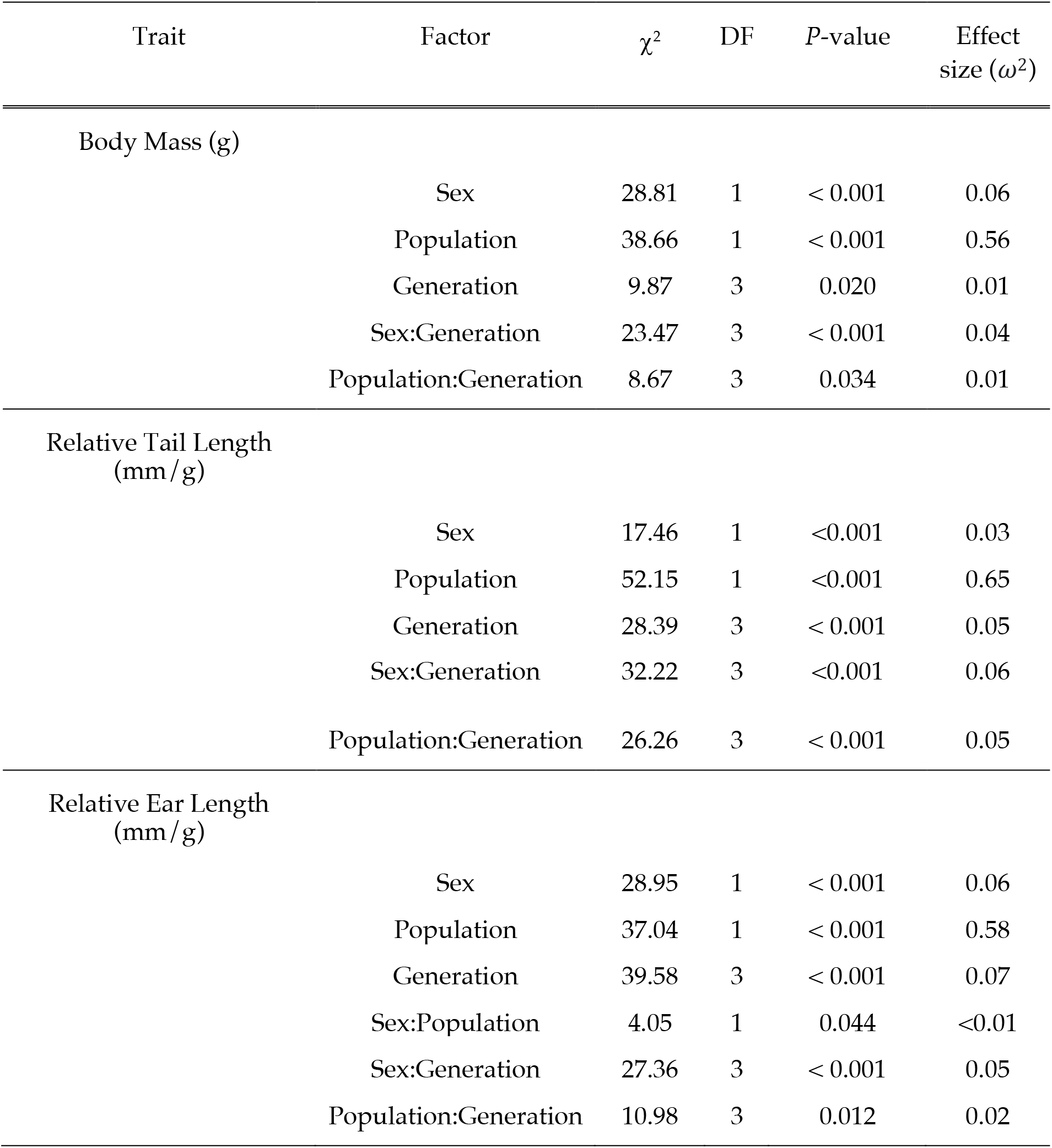
Results of linear mixed models investigating the effects of sex, population, generation, and their interaction on body mass and extremity length in house mice. Results of analysis are plotted in Figure S2.

## Supplemental Figures

**Figure S1.**
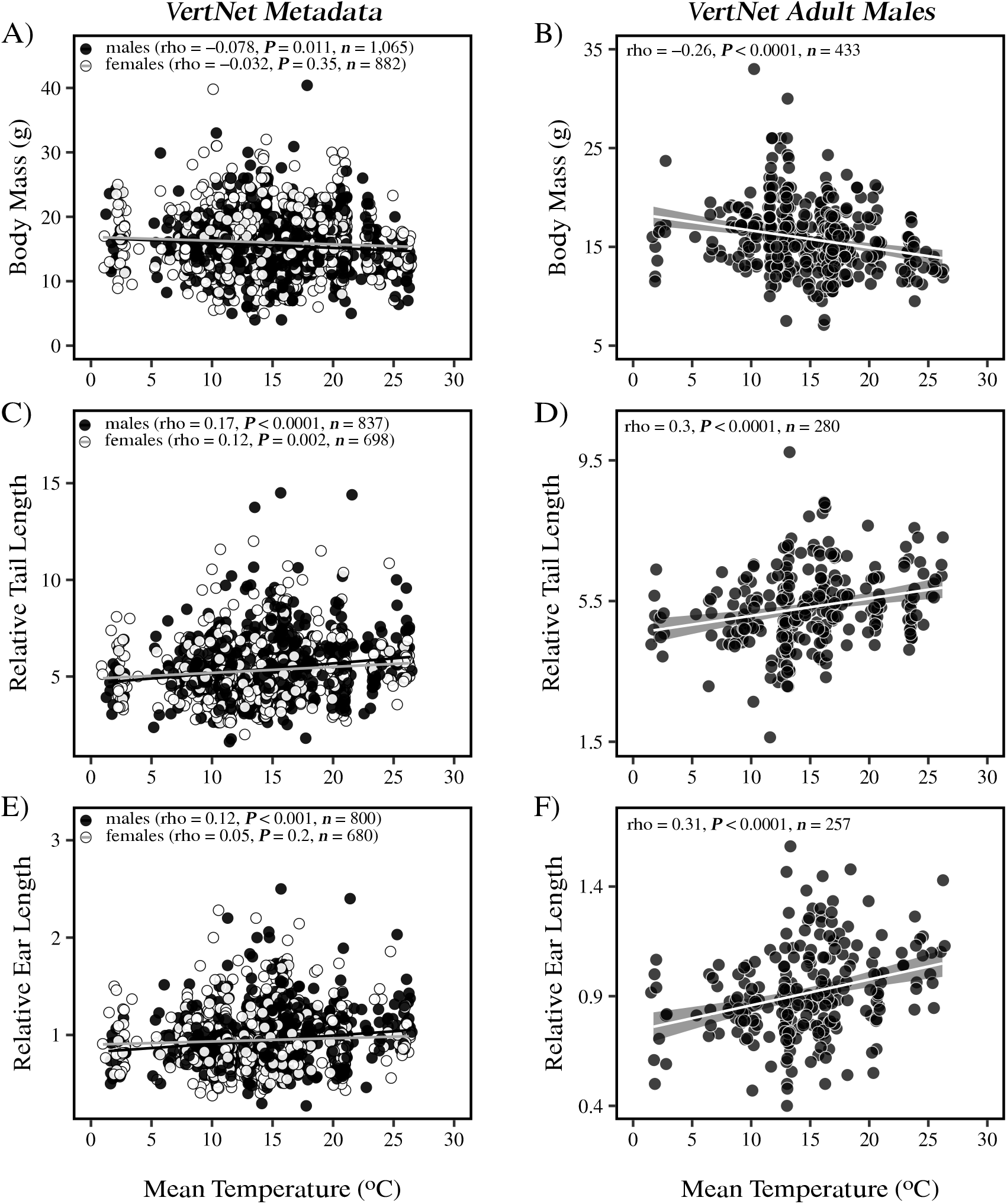
Bergmann’s rule and Allen’s rule in house mice from North and South America. Associations between body mass (A-B), tail length (C-D), ear length (E-F), and mean annual temperature (°C) across wild-caught North and South American house mice. Tail length and ear length are plotted relative to body mass for each individual. Individuals are represented as individual points, with males depicted in black and females depicted in white. Results from Spearman correlations are presented in each plot, along with sample sizes. For clarity, standard error shading is omitted from linear regression lines associated with the VertNet Metadata (panels A, C, and E).

**Figure S2.**
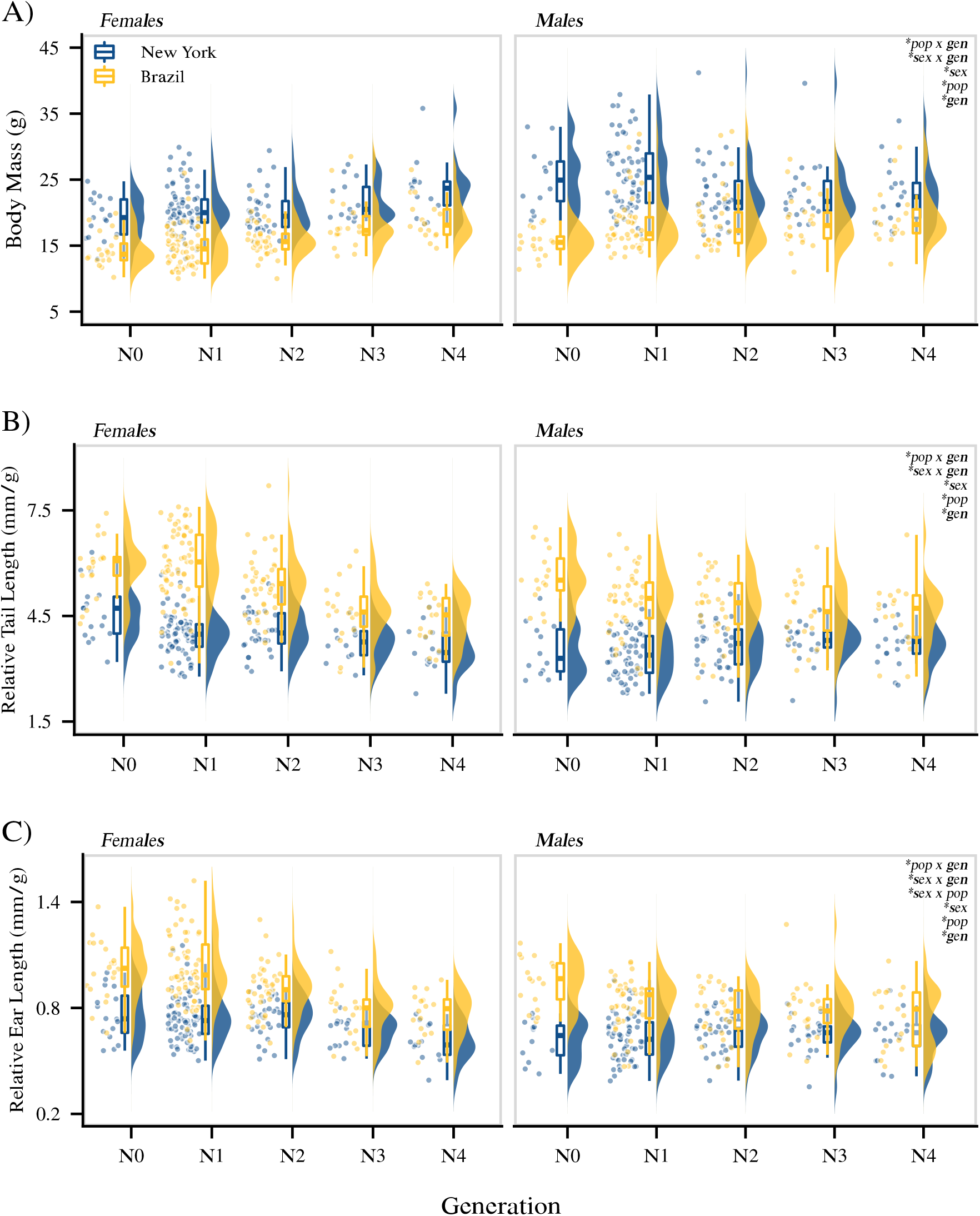
Body mass and extremity length differences among populations persist over generations in a common lab environment. Differences in body mass (A), tail length (B), and ear length (C) between New York mice (blue) and Brazil mice (gold) across generations. Tail length and ear length are plotted relative to body mass for each individual. Population-level data are depicted as boxplots overlayed on density plots, with boxplot vertical lines denoting 1.5x the inerquartile range. Individuals are represented as individual points. Results from linear models are presented in the upper right corner of each panel (\**P*<0.05; Table S2). Abbreviations: N0 = generation 0 (wild mice); N1 = generation 1; N2-N4 = generations 2-4. Sample sizes: (A) Females (New York: *n* = 146; Brazil: *n* = 157); Males (New York: *n* = 140; Brazil: *n* = 133); (B) Females (New York: *n* = 145; Brazil: *n* = 156); Males (New York: *n* = 140; Brazil: *n* = 125); (C) Females (New York: *n* = 145; Brazil: *n* = 156); Males (New York: *n* = 139; Brazil: *n* = 128).

**Figure S3.**
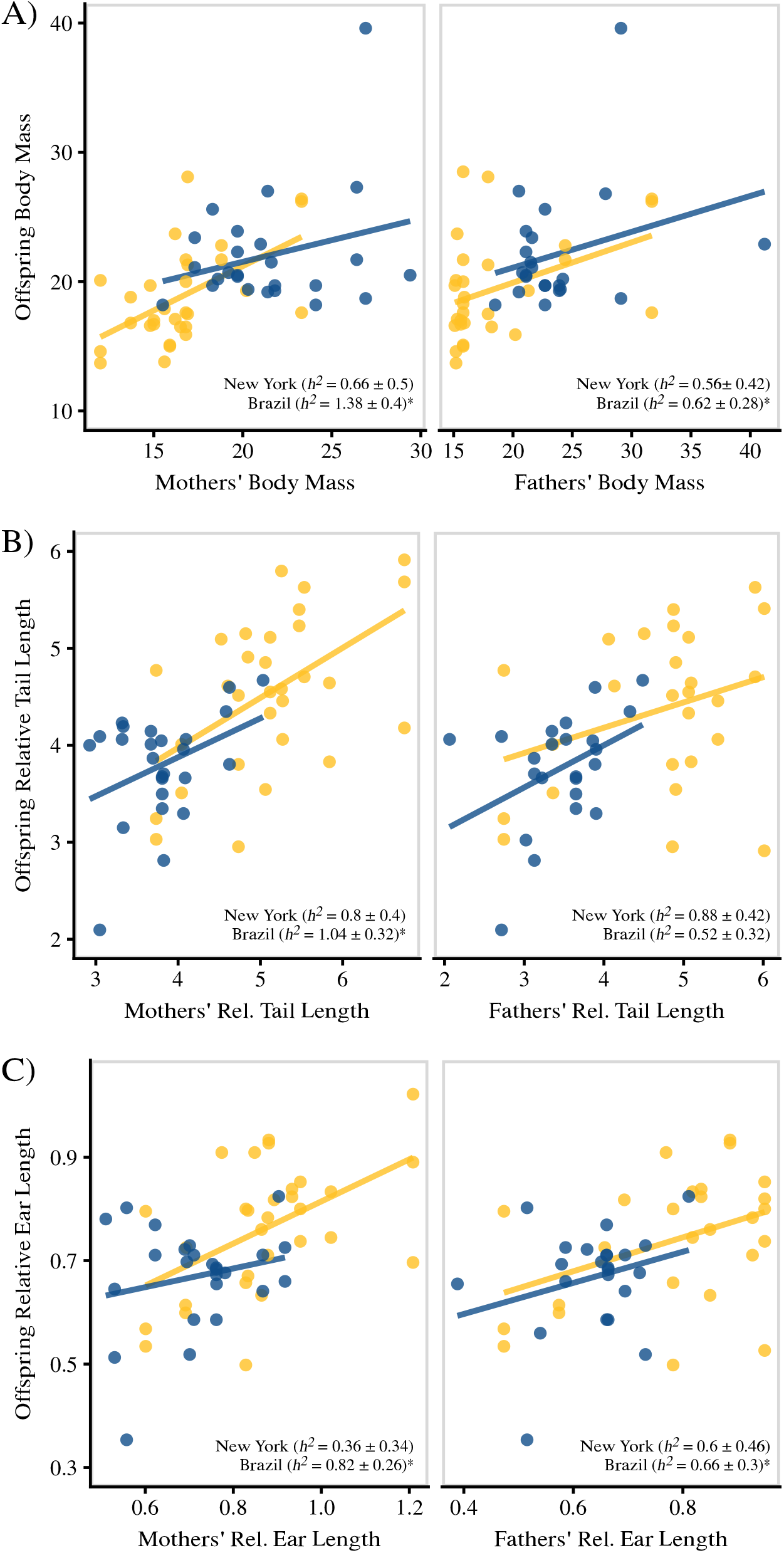
Maternal- and paternal-offspring heritability (*h*^2^) estimates for body mass and extremity length in New York and Brazil house mice. Maternal-offspring and paternal-offspring regressions were performed on body mass (A), relative tail length (B), and relative ear length (C). Heritabilities (*h*^2^ ± standard error) were estimated as twice the slope and standard error of single-parent offspring regressions. Heritability values higher than one are the result of doubling statistics with large standard errors. Significance of each regression were assessed with ANOVAs and are represented with an asterisk. Sample sizes: New York: *n* = 25; Brazil: *n* = 30.

**Figure S4.**
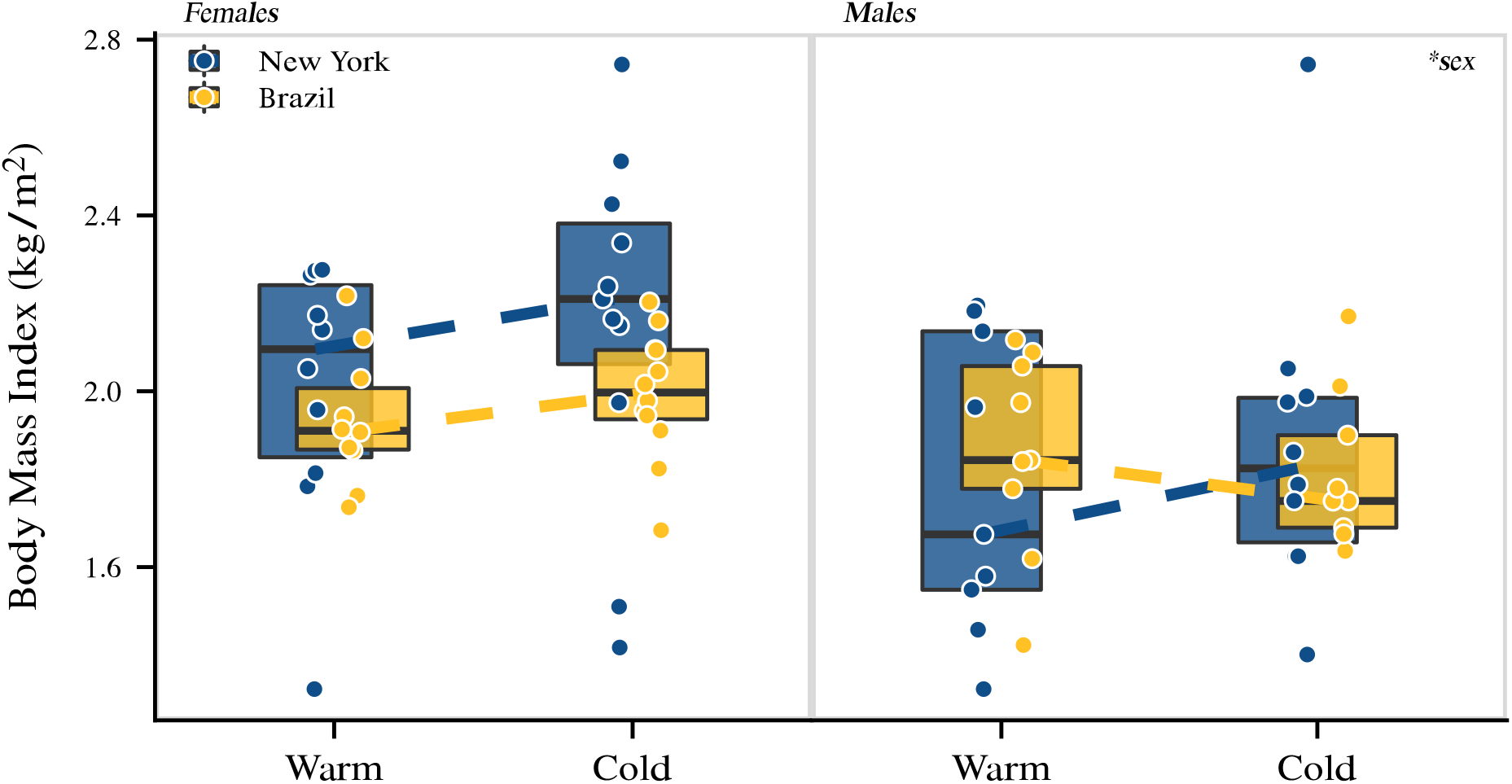
No genetic differences or plasticity in body mass index (BMI) among New York mice and Brazil mice. Individuals are represented as individual points (New York: *n* = 20 per treatment; Brazil: *n* = 20 per treatment), and boxplots indicate the 25th, median, and 75th quartiles. Dashed lines connect median values of each genotype across warm and cold environments. The same individuals depicted here are also depicted in Figure 3.

## References

Alhajeri, B. H., Y. Fourcade, N. S. Upham, and H. Alhaddad. 2020. A global test of Allen’s rule in rodents. Global Ecology and Biogeography 29:2248–2260.

Alhajeri, B. H., and S. J. Steppan. 2016. Association between climate and body size in rodents: A phylogenetic test of Bergmann’s rule. Mammalian Biology 81:219–225.

Al-Hilli, F., and E. Wright. 1983. The effects of changes in the environmental temperature on the growth of bone in the mouse. Radiological and morphological study. British Journal of Experimental Pathology 64:43.

Alho, J., G. Herczeg, A. Laugen, K. Räsänen, A. Laurila, and J. Merilä. 2011. Allen’s rule revisited: Quantitative genetics of extremity length in the common frog along a latitudinal gradient. Journal of Evolutionary Biology 24:59–70.

Allaire, J., Y. Xie, J. McPherson, J. Luraschi, K. Ushey, A. Atkins, H. Wickham, et al. 2021. Rmarkdown: Dynamic documents for R.

Allen, J. A. 1877. The influence of physical conditions in the genesis of species. Radical Review 1:108–140.

Alroy, J. 2019. Small mammals have big tails in the tropics. Global Ecology and Biogeography 28:1042–1050.

Andrew, S., L. Hurley, M. Mariette, and S. Griffith. 2017. Higher temperatures during development reduce body size in the zebra finch in the laboratory and in the wild. Journal of Evolutionary Biology 30:2156–2164.

Ashoub, M. E.-R. 1958. Effect of two extreme temperatures on growth and tail-length of mice. Nature 181:284–284.

Ashton, K. G. 2002. Patterns of within-species body size variation of birds: Strong evidence for Bergmann’s rule. Global Ecology and Biogeography 11:505–523.

Ashton, K. G., M. C. Tracy, and A. de Queiroz. 2000. Is Bergmann’s rule valid for mammals? The American Naturalist 156:390–415.

Baldwin, J. M. 1896. A new factor in evolution. The American Naturalist 30:441–451.

Barnett, S. 1965. Genotype and environment in tail length in mice. Quarterly Journal of Experimental Physiology and Cognate Medical Sciences: Translation and Integration 50:417–429.

Barnett, S., and R. Dickson. 1984. Changes among wild house mice (Mus musculus) bred for ten generations in a cold environment, and their evolutionary implications. Journal of Zoology 203:163–180.

Bates, D., M. Mächler, B. Bolker, and S. Walker. 2015. Fitting linear mixed-effects models using lme4. Journal of Statistical Software 67:1–48.

Ben-Shachar, M. S., D. Lüdecke, and D. Makowski. 2020. effectsize: Estimation of effect size indices and standardized parameters. Journal of Open Source Software 5:2815.

Bergmann, C. 1847. Über die verhältnisse der wärmeökonomie der thiere zu ihrer grösse. Gottinger Studien 3:595–708.

Betti, L., S. J. Lycett, N. von Cramon-Taubadel, and O. M. Pearson. 2015. Are human hands and feet affected by climate? A test of Allen’s rule. American Journal of Physical Anthropology 158:132–140.

Blackburn, T. M., and B. A. Hawkins. 2004. Bergmann’s rule and the mammal fauna of northern North America. Ecography 27:715–724.

Brown, J. H., and A. K. Lee. 1969. Bergmann’s rule and climatic adaptation in woodrats (Neotoma). Evolution 329–338.

Burness, G., J. R. Huard, E. Malcolm, and G. J. Tattersall. 2013. Post-hatch heat warms adult beaks: Irreversible physiological plasticity in Japanese quail. Proceedings of the Royal Society B: Biological Sciences 280:20131436.

Cardilini, A. P., K. L. Buchanan, C. D. Sherman, P. Cassey, and M. R. Symonds. 2016. Tests of ecogeographical relationships in a non-native species: What rules avian morphology? Oecologia 181:783–793.

Cavicchi, S., D. Guerra, G. Giorgi, and C. Pezzoli. 1985. Temperature-related divergence in experimental populations of Drosophila melanogaster. I. Genetic and developmental basis of wing size and shape variation. Genetics 109:665–689.

Chevillard, L., R. Portet, and C. M. 1963. Growth rate of rats born and reared at 5 and 30 c. Federation Proceedings 22:699–703.

Conover, D. O., and E. T. Schultz. 1995. Phenotypic similarity and the evolutionary significance of countergradient variation. Trends in Ecology & Evolution 10:248–252.

Constable, H., R. Guralnick, J. Wieczorek, C. Spencer, A. T. Peterson, V. S. Committee, and others. 2010. VertNet: A new model for biodiversity data sharing. PLoS Biology 8:e1000309.

Coyne, J. A., and E. Beecham. 1987. Heritability of two morphological characters within and among natural populations of Drosophila melanogaster. Genetics 117:727–737.

Danner, R. M., and R. Greenberg. 2015. A critical season approach to Allen’s rule: Bill size declines with winter temperature in a cold temperate environment. Journal of Biogeography 42:114–120.

Des Marais, D. L., K. M. Hernandez, and T. E. Juenger. 2013. Genotype-by-environment interaction and plasticity: Exploring genomic responses of plants to the abiotic environment. Annual Review of Ecology, Evolution, and Systematics 44:5–29.

Endler, J. A. 1977. Geographic variation, speciation, and clines. Princeton University Press.

Falconer, D. S., and T. F. C. Mackay. 1996. Introduction to quantitative genetics. Pearson.

Ferris, K. G., A. S. Chavez, T. A. Suzuki, E. J. Beckman, M. Phifer-Rixey, K. Bi, and M. W. Nachman. 2021. The genomics of rapid climatic adaptation and parallel evolution in North American house mice. PLoS Genetics 17:e1009495.

Fick, S. E., and R. J. Hijmans. 2017. WorldClim 2: New 1-km spatial resolution climate surfaces for global land areas. International journal of climatology 37:4302–4315.

Fooden, J., and G. H. Albrecht. 1999. Tail-length evolution in fascicularis-group macaques (Cercopithecidae: macaca). International Journal of Primatology 20:431–440.

Foster, F., and M. Collard. 2013. A reassessment of Bergmann’s rule in modern humans. PloS One 8:e72269.

Fox, J., and S. Weisberg. 2019. An R companion to applied regression (Third.). Sage, Thousand Oaks CA.

Freckleton, R. P., P. H. Harvey, and M. Pagel. 2003. Bergmann’s rule and body size in mammals. The American Naturalist 161:821–825.

Friedman, N. R., L. Harmáčková, E. P. Economo, and V. Remeš. 2017. Smaller beaks for colder winters: Thermoregulation drives beak size evolution in Australasian songbirds. Evolution 71:2120–2129.

Geist, V. 1987. Bergmann’s rule is invalid. Canadian Journal of Zoology 65:1035–1038.

Ghalambor, C. K., J. K. McKay, S. P. Carroll, and D. N. Reznick. 2007. Adaptive versus non-adaptive phenotypic plasticity and the potential for contemporary adaptation in new environments. Functional Ecology 21:394–407.

Gilchrist, G. W., R. B. Huey, J. Balanyà, M. Pascual, and L. Serra. 2004. A time series of evolution in action: A latitudinal cline in wing size in South American Drosophila subobscura. Evolution 58:768–780.

Gilchrist, G. W., R. B. Huey, and L. Serra. 2001. Rapid evolution of wing size clines in Drosophila subobscura. Genetica 273–286.

Gillespie, J. H., and M. Turelli. 1989. Genotype-environment interactions and the maintenance of polygenic variation. Genetics 121:129–138.

Gohli, J., and K. L. Voje. 2016. An interspecific assessment of Bergmann’s rule in 22 mammalian families. BMC Evolutionary Biology 16:1–12.

Gomulkiewicz, R., and M. Kirkpatrick. 1992. Quantitative genetics and the evolution of reaction norms. Evolution 46:390–411.

Gordon, C. 2012. Thermal physiology of laboratory mice: Defining thermoneutrality. Journal of Thermal Biology 37:654–685.

Griffing, J. P. 1974. Body measurements of black-tailed jackrabbits of southeastern New Mexico with implications of Allen’s rule. Journal of Mammalogy 55:674–678.

Harland, S. 1960. Effect of temperature on growth in weight and tail-length of inbred and hybrid mice. Nature 186:446.

Harpak, A., and M. Przeworski. 2021. The evolution of group differences in changing environments. PLoS Biology 19:e3001072.

Huey, R. B., G. W. Gilchrist, M. L. Carlson, D. Berrigan, and L. Serra. 2000. Rapid evolution of a geographic cline in size in an introduced fly. Science 287:308–309.

Husby, A., S. M. Hille, and M. E. Visser. 2011. Testing mechanisms of Bergmann’s rule: Phenotypic decline but no genetic change in body size in three passerine bird populations. The American Naturalist 178:202–213.

Huxley, J. S. 1939. Clines: An auxiliary method in taxonomy. Bijdragen tot de Dierkunde 27:491–520.

James, A. C., R. Azevedo, and L. Partridge. 1995. Cellular basis and developmental timing in a size cline of Drosophila melanogaster. Genetics 140:659–666.

James, F. C. 1970. Geographic size variation in birds and its relationship to climate. Ecology 51:365–390.

James, F. C. 1983. Environmental component of morphological differentiation in birds. Science 221:184–186.

Johnston, R. F., and R. K. Selander. 1964. House sparrows: Rapid evolution of races in North America. Science 144:548–550.

Johnston, R. F., and R. K. Selander. 1971. Evolution in the house sparrow. II. Adaptive differentiation in North American populations. Evolution 1–28.

Laiolo, P., and A. Rolando. 2001. Ecogeographic correlates of morphometric variation in the red-billed chough Pyrrhocorax pyrrhocorax and the alpine chough Pyrrhocorax graculus. Ibis 143:602–616.

Land, J. V. ‘ t., P. V. Putten, and W. V. Delden. 1999. Latitudinal variation in wild populations of Drosophila melanogaster: Heritabilities and reaction norms. Journal of Evolutionary Biology 12:222–232.

Lüdecke, D., M. S. Ben-Shachar, I. Patil, P. Waggoner, and D. Makowski. 2021. Assessment, testing and comparison of statistical models using R. Journal of Open Source Software 6:3112.

Lynch, C. B. 1992. Clinal variation in cold adaptation in Mus domesticus: Verification of predictions from laboratory populations. The American Naturalist 139:1219–1236.

Lynch, M., B. Walsh, and others. 1998. Genetics and analysis of quantitative traits. Sinauer Sunderland, MA.

Mayr, E. 1956. Geographical character gradients and climatic adaptation. Evolution 10:105–108.

McNab, B. K. 1971. On the ecological significance of Bergmann’s rule. Ecology 52:845–854.

Meiri, S., and T. Dayan. 2003. On the validity of Bergmann’s rule. Journal of Biogeography 30:331–351.

Millien, V., S. Kathleen Lyons, L. Olson, F. A. Smith, A. B. Wilson, and Y. Yom-Tov. 2006. Ecotypic variation in the context of global climate change: Revisiting the rules. Ecology Letters 9:853–869.

Mincer, S. T., and G. A. Russo. 2020. Substrate use drives the macroevolution of mammalian tail length diversity. Proceedings of the Royal Society B 287:20192885.

Noel, J. F., and E. Wright. 1970. The effect of environmental temperature on the growth of vertebrae in the tail of the mouse. Development 24:405–410.

Nudds, R., and S. Oswald. 2007. An interspecific test of Allen’s rule: Evolutionary implications for endothermic species. Evolution: International Journal of Organic Evolution 61:2839–2848.

Ogle, C., and C. Mills. 1933. Animal adaptation to environmental temperature conditions. American Journal of Physiology-Legacy Content 103:606–612.

Olejnik, S., and J. Algina. 2003. Generalized eta and omega squared statistics: Measures of effect size for some common research designs. Psychological methods 8:434.

Olson, V. A., R. G. Davies, C. D. L. Orme, G. H. Thomas, S. Meiri, T. M. Blackburn, K. J. Gaston, et al. 2009. Global biogeography and ecology of body size in birds. Ecology Letters 12:249–259.

Ozgul, A., D. Z. Childs, M. K. Oli, K. B. Armitage, D. T. Blumstein, L. E. Olson, S. Tuljapurkar, et al. 2010. Coupled dynamics of body mass and population growth in response to environmental change. Nature 466:482–485.

Ozgul, A., S. Tuljapurkar, T. G. Benton, J. M. Pemberton, T. H. Clutton-Brock, and T. Coulson. 2009. The dynamics of phenotypic change and the shrinking sheep of St. kilda. Science 325:464–467.

Partridge, L., B. Barrie, K. Fowler, and V. French. 1994. Evolution and development of body size and cell size in Drosophila melanogaster in response to temperature. Evolution 48:1269–1276.

Phifer-Rixey, M., K. Bi, K. G. Ferris, M. J. Sheehan, D. Lin, K. L. Mack, S. M. Keeble, et al. 2018. The genomic basis of environmental adaptation in house mice. PLoS Genetics 14:e1007672.

Phifer-Rixey, M., and M. W. Nachman. 2015. The natural history of model organisms: Insights into mammalian biology from the wild house mouse Mus musculus. Elife 4:e05959.

Price, T. D., A. Qvarnström, and D. E. Irwin. 2003. The role of phenotypic plasticity in driving genetic evolution. Proceedings of the Royal Society of London. Series B: Biological Sciences 270:1433–1440.

Riemer, K., R. P. Guralnick, and E. P. White. 2018. No general relationship between mass and temperature in endothermic species. Elife 7:e27166.

Romano, A., R. Séchaud, and A. Roulin. 2020. Geographical variation in bill size provides evidence for Allen’s rule in a cosmopolitan raptor. Global Ecology and Biogeography 29:65–75.

Ruff, C. 2002. Variation in human body size and shape. Annual Review of Anthropology 31:211–232.

Ruff, C. B. 1994. Morphological adaptation to climate in modern and fossil hominids. American Journal of Physical Anthropology 37:65–107.

Rutledge, J. J., E. Eisen, and J. Legates. 1973. An experimental evaluation of genetic correlation. Genetics 75:709–726.

Scholander, P. F. 1955. Evolution of climatic adaptation in homeotherms. Evolution 15–26.

Serrat, M. A. 2013. Allen’s rule revisited: Temperature influences bone elongation during a critical period of postnatal development. The Anatomical Record 296:1534–1545.

Serrat, M. A. 2014. Environmental temperature impact on bone and cartilage growth. Comprehensive Physiology 4:621–655.

Serrat, M. A., D. King, and C. O. Lovejoy. 2008. Temperature regulates limb length in homeotherms by directly modulating cartilage growth. Proceedings of the National Academy of Sciences 105:19348–19353.

Sumner, F. B. 1909. Some effects of external conditions upon the white mouse. The Journal of Experimental Zoology 7:97–155.

Sumner, F. B. 1915. Some studies of environmental influence, heredity, correlation and growth, in the white mouse. The Journal of Experimental Zoology 18:325.

Suzuki, T. A., F. M. Martins, Phifer-Rixey Megan, and M. W. Nachman. 2020. The gut microbiota and Bergmann’s rule in wild house mice. Molecular Ecology 29:2300–2311.

Symonds, M. R., and G. J. Tattersall. 2010. Geographical variation in bill size across bird species provides evidence for Allen’s rule. The American Naturalist 176:188–197.

Tattersall, G. J., B. Arnaout, and M. R. Symonds. 2017. The evolution of the avian bill as a thermoregulatory organ. Biological Reviews 92:1630–1656.

Teplitsky, C., J. A. Mills, J. S. Alho, J. W. Yarrall, and J. Merilä. 2008. Bergmann’s rule and climate change revisited: Disentangling environmental and genetic responses in a wild bird population. Proceedings of the National Academy of Sciences 105:13492–13496.

Thorington Jr, R. W. 1970. Lability of tail length of the white-footed mouse, Peromyscus leucopus noveboracensis. Journal of Mammalogy 51:52–59.

Tomlinson, S., and P. C. Withers. 2009. Biogeographical effects on body mass of native Australian and introduced mice, Pseudomys hermannsburgensis and Mus domesticus: An inquiry into Bergmann’s rule. Australian Journal of Zoology 56:423–430.

Via, S., and R. Lande. 1985. Genotype-environment interaction and the evolution of phenotypic plasticity. Evolution 39:505–522.

Weaver, M. E., and D. L. Ingram. 1969. Morphological changes in swine associated with environmental temperature. Ecology 50:710–713.

West-Eberhard, M. J. 2003. Developmental plasticity and evolution. Oxford University Press.

Wickham, H., M. Averick, J. Bryan, W. Chang, L. D. McGowan, R. François, G. Grolemund, et al. 2019. Welcome to the tidyverse. Journal of Open Source Software 4:1686.

Yang, J., B. Benyamin, B. P. McEvoy, S. Gordon, A. K. Henders, D. R. Nyholt, P. A. Madden, et al. 2010. Common SNPs explain a large proportion of the heritability for human height. Nature genetics 42:565–569.

Yom-Tov, Y., and H. Nix. 1986. Climatological correlates for body size of five species of Australian mammals. Biological Journal of the Linnean Society 29:245–262.

